# Predictive Maps in Rats and Humans for Spatial Navigation

**DOI:** 10.1101/2020.09.26.314815

**Authors:** William de Cothi, Nils Nyberg, Eva-Maria Griesbauer, Carole Ghanamé, Fiona Zisch, Julie M. Lefort, Lydia Fletcher, Coco Newton, Sophie Renaudineau, Daniel Bendor, Roddy Grieves, Éléonore Duvelle, Caswell Barry, Hugo J. Spiers

## Abstract

Much of our understanding of navigation comes from the study of individual species, often with specific tasks tailored to those species. Here, we provide a novel experimental and analytic framework, integrating across humans, rats and simulated reinforcement learning (RL) agents to interrogate the dynamics of behaviour during spatial navigation. We developed a novel open-field navigation task (ʻTartarus Maze’) requiring dynamic adaptation (shortcuts and detours) to frequently changing obstructions in the path to a hidden goal. Humans and rats were remarkably similar in their trajectories. Both species showed the greatest similarity to RL agents utilising a ʻsuccessor representation’, which creates a predictive map. Humans also displayed trajectory features similar to model-based RL agents, which implemented an optimal tree-search planning procedure. Our results help refine models seeking to explain mammalian navigation in dynamic environments, and highlight the utility of modelling the behaviour of different species to uncover the shared mechanisms that support behaviour.

## Introduction

Adapting to change is fundamental for survival. Adapting to changes in the structure of the environment has been studied in a huge diversity of psychological experiments in humans^1^, but also more ethologically in a remarkable range of different species^2^. One challenge that unites all motile animals on our planet is spatial navigation. In particular, prime examples are finding a new path when a familiar route is blocked and exploiting a novel shortcut. Efficient detours and shortcuts are considered the hallmarks of a cognitive map - an internal representation of the environment that enables novel inferences to guide behaviour^3–6^.

Both rodents and humans can show an impressive capacity to identify shortcuts and take optimal detours^5, 7–16^. However, not all studies report successful adaptive behaviour^17^. Rats often require multiple exposures to a set of paths before they are able to shift towards an optimal detour ^9^, and may fail to select an optimal shortcut from a set of novel paths ^18^. Humans too can be poor at judging the directions between locations in walled mazes, hindering the capacity to identify shortcuts^19, 20^.

Much of the research into navigation implicitly assumes that rodents and humans navigate in a fundamentally similar way^4, 21^ and this has been used to support the integration of insights across both species^1, 5, 22–25^. In mammals, the hippocampus is thought to form a cognitive map^24^, evidenced by spatial-tuned cells (such as ʻplace cells’) in the hippocampal formation of rodents and humans^26–28^. However, despite the wide array of human and rodent research, few experiments have sought to compare rodents and humans on a directly homologous task. Understanding the similarities and differences of these two species on the same task would be useful for allowing the better integration of findings from different methods, such as combining data from neuroimaging in humans with neural recordings and disruption methods in rodents^29–31^. Moreover, such integration could potentially benefit the translation of assessments in rodents to assessments for clinical trials in humans, for example where tests of spatial navigation may be important for the early detection of Alzheimer’s disease^32–34^.

When considering how humans and rodents might differ during navigation, differences in sensory perception are important. Whilst humans have binocular vision, they may differ in olfaction^35^ and lack the tactility of whiskers. Meanwhile, rodents have a larger visual field of view, lower visual acuity and can move their eyes independently^36^. In terms of neuroanatomy, the prefrontal cortical regions associated with spatial planning differ greatly between rodents and primates^37, 38^; while the hippocampus and surrounding structures associated with spatial representations are relatively similar^39^. Given these similarities and differences, it is possible that rodents and humans navigate in a similar fashion or show pronounced differences in certain situations. Understanding such patterns in behaviour is important not only for understanding navigation, but how the behaviour of different species is inter-related and may have emerged through evolutionary pressure.

One approach for identifying potential cross-species mechanisms underlying goal-directed behaviour is through comparison with reinforcement learning models^40–45^. Reinforcement learning (RL) is an area of machine learning that addresses the theoretical problem of how a learner and decision maker, called an agent, should act in an environment in order to achieve a certain goal - for which it earns rewards. Specifically, the agent is not told which actions it should take, but instead must learn the actions that maximise its expected future rewards - known as value. Such RL models can be used to examine how rapid learning and control can be developed in artificial systems, outcompeting human performance^40, 46–48^, or used for comparison to patterns seen in animals or human behaviour^45, 49–52^.

Solutions to reinforcement learning problems have traditionally been divided into two categories: model-based methods that afford the agent a model of the environment, used to decide actions via a planning procedure^53^, and model-free methods that learn from experience which actions lead to the most rewarding future^54, 55^. Provided that the model implemented in a model-based algorithm contains an accurate depiction of the environment, model-based methods are typically able to respond quickly and optimally to environmental perturbations. However, the planning procedure - for example a tree search^47^ - required to successfully exploit the model brings with it computational complexity and overhead, particularly in large state spaces with deep transition structures such as navigating a city.

In contrast to model-based methods, model-free methods are generally more simple and computationally inexpensive through a reliance on temporal-difference learning rules^54^, however this comes with a reduced flexibility to environmental changes. As such, model-free mechanisms are often associated with the formation of habits^56, 57^. To achieve their simplicity, model-free methods typically learn by directly estimating the value of taking a particular action in a particular state. This makes it easy to then compare the values of different actions available to the agent, without the need to know how the states are interconnected.

Whilst model-free and model-based methods appear to function at opposite ends of an algorithmic spectrum, intermediary methods do exist. One such algorithm that has increased in application recently is the successor representation^58^ (SR). The SR somewhat combines parts of model-free and model-based learning^59, 60^ by using experience to learn a predictive map between the states in an environment. This predictive map can be readily combined with a separately learned reward associated with each state, in order to explicitly compute value. Thus the SR negates the need for a complicated planning procedure in order to use the predictive map to guide action selection.

The SR has been able to provide a good account of behaviour and hippocampal representations in humans^61–65^ and rodents^50, 66, 67^. The tasks often used to draw these comparisons with RL agents typically focus on small state-spaces, with 2-step transition structures – as such the extent of planning often requires one or two actions. Furthermore, due to the conceptual nature of the underlying task space, translational research usually requires differing sensory implementations for humans^68^ and rodents^69^.

Here, we created a configurable open-field maze with a layout of barriers that reconfigured after a set of trials (Tartarus Maze). We tested the navigation of rats in a physical instantiation of the maze, humans via an immersive head-mounted display virtual environment and RL agents in a simulation. Using a range of analytic methods we probed how rat and human spatial behaviours compare to each other and to model-free, model-based and SR reinforcement learners. We found a strong similarity in the occupancy patterns of rats and humans. Both rats and humans showed the greatest likelihood and trajectory similarity to SR based RL agents, with humans also displaying trajectory features similar to model-based RL agents implementing an optimal planning procedure in early trials on a new maze configuration.

## Results

Navigation was tested in a large square environment with a fixed hidden goal location and a prominent directional black wall cue in one direction (Fig. 1, Video S1 - rats, and Video S2 - humans). The maze was divided in a 10×10 grid of moveable sections that could either be removed, leaving impassable gaps to force detour taking, or added, creating shortcuts. The speed and size of the humans in the virtual environment was set to match a rat travelling at 20cm/s. During training, all 10×10 maze modules were present and the rats and humans were trained to reach the goal within a 45s time limit, (Fig 1A), while RL agents were initialised with the optimal value function. During the testing phase of the experiment, maze modules were removed to make specific maze configurations that block the direct route to the goal (Fig 1B). Humans (n=18), rats (n=9) and agents were tested on the same sequence of 25 maze configurations each with 10 trials in which a set of defined starting locations were selected to optimally probe navigation (Fig 1C). See Methods for details.

**Figure 1:**
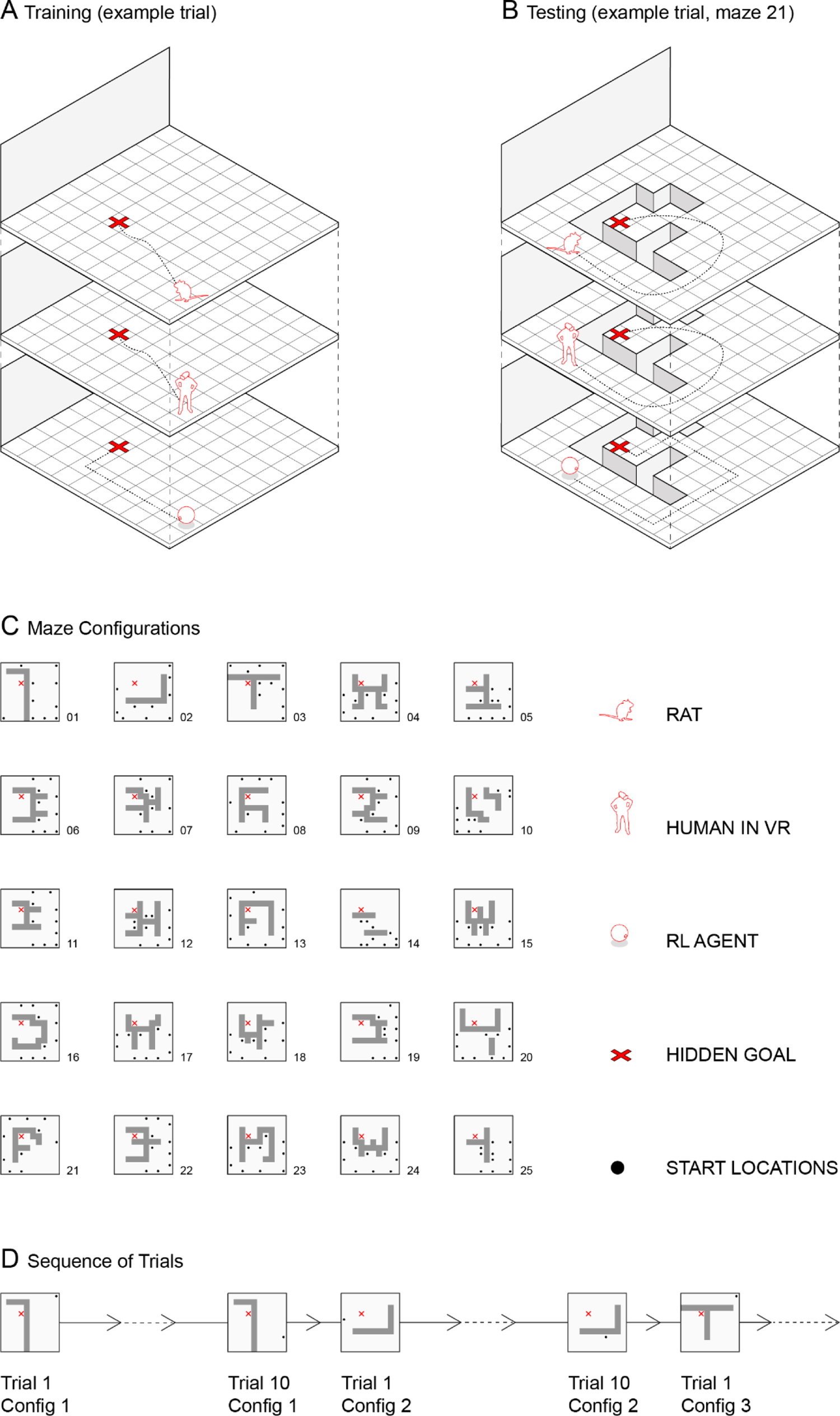
The Tartarus Maze. **(A)** Schematic of the maze composed of 10×10 units for humans, rats and RL agents. For rats each unit (20×20cm) had a port to potentially deliver a chocolate milk reward after the rat waited 5 sec at the goal (See Video S1). For humans each unit could be associated with a hidden gold star linked to financial reward, which appeared after waiting 5 sec at the goal location. Gaps between units were not visible to humans to avoid counting distance to the hidden goal (see Video S2). Example trial shows one of the possible pseudo-random starting locations on the edge of the maze. **(B)** After training, flexible navigation was tested by removing units from the maze to create maze configurations with gaps between traversable surfaces. Example from maze configuration 21 is shown with one of the 10 starting locations tested. Each configuration was tested for 10 trials, with each trial having a different starting location. **(C**) Sequence of 25 maze configurations used. **(D)** Illustration of the trial sequence, highlighting the transition in layout every 10 trials across the 250 trials tested in the 25 maze configurations.

### Behavioural performance is relatively similar between rats and humans

We first asked how well humans (Fig 2A) and rodents (Fig 2B) were able to complete the task during the test sessions. As expected, repeated exposure over trials to a new maze configuration corresponded to a general increase in the ability of both the humans and rats to navigate to the goal within the 45s time limit (Fig 2C; first 5 trials vs last 5 trials: humans t(17)=6.3, *p<*.001; rats t(8)=4.0, *p=* .004). Humans were also generally better than the rats at finding the goal across the 25 maze configurations (Fig 2D; humans vs rats: t(25)=3.0, *p=*.006). There were 3 maze configurations in which rats outperformed humans (2, 10 and 19). We saw a strong correlation between the occupancy of the rats and human participants (occupancy correlation, humans vs rats: ρ=.67), in particular towards the later trials when both were better at navigating to the goal (Fig 2E; occupancy correlations for first 5 trials vs last 5 trials: t(8)=3.2, *p=*.013). The routes used were also more efficient (Fig 2E; deviation from optimal path, first 5 trials vs last 5 trials: human t(17)=-5.0, *p<*.001; rats t(8)=-4.0, *p=*.004) with humans generally choosing more optimal routes than the rats (deviation from optimal path humans vs rats: t(25)=-8.2, *p*<.001). Inspection of trajectories showed that in some cases near optimal paths could be observed in both species even on the first trial of a maze configuration (see Video S1&2).

**Figure 2:**
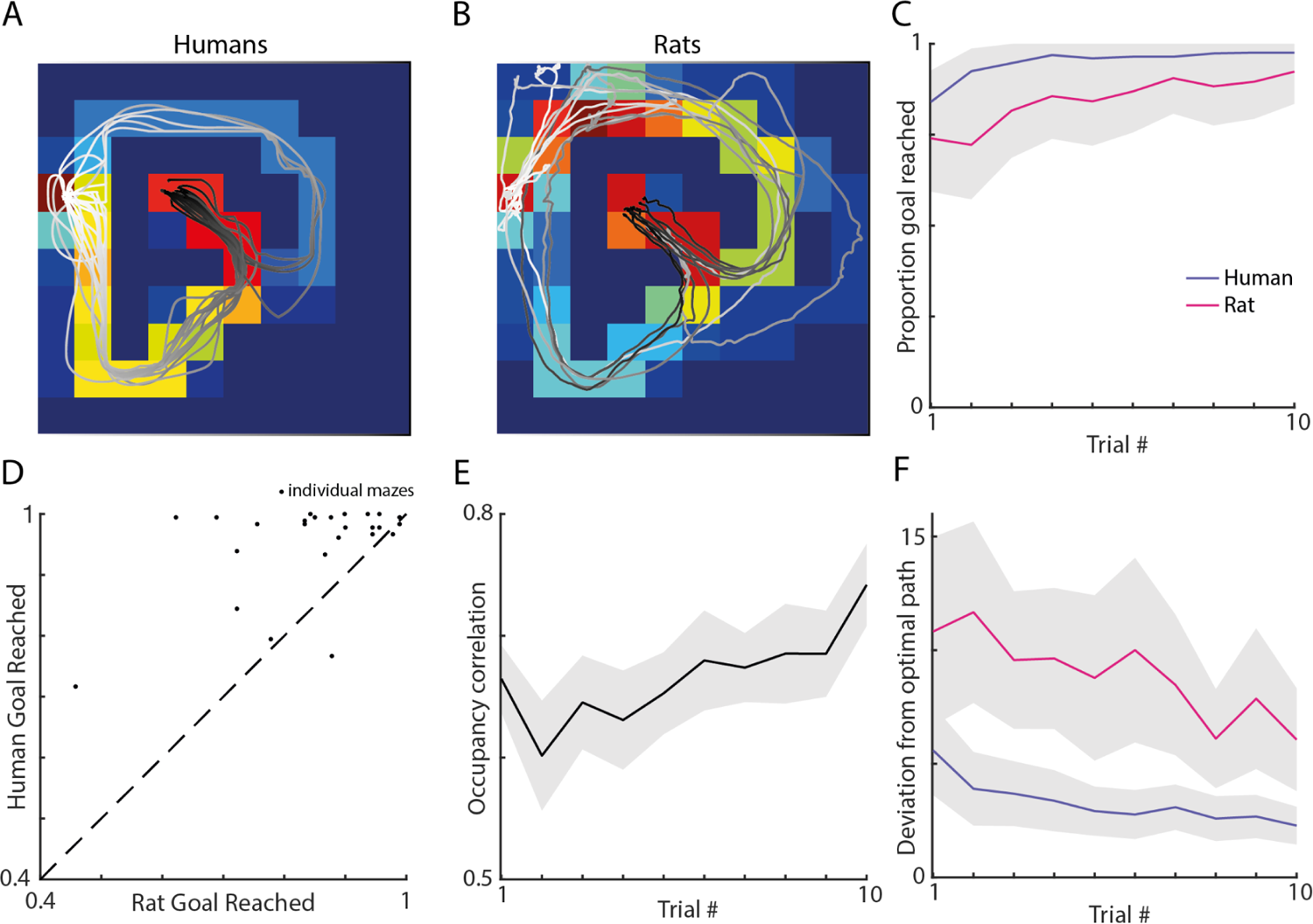
Humans and rats were able to successfully navigate to the hidden goal and generally did so using similar routes. Examples of the human **(A)** and rat **(B)** trajectories overlaying occupancy maps for a given trial. The white-black colour gradient shows the beginning-end of each trajectory. **(C)** Proportion of trials where the goal was reached averaged over all configurations as a function of trial number, for rats (red) or humans (green). Grey areas indicate standard error from the mean. **(D)** Proportion of goals reached by humans and rats during the time limit. The rats outperformed humans on a total of three maze configurations (2,10,19), which was most pronounced for configuration 2. Configuration 1 was the configuration both species performed most inaccurately on. **(E)** Correlation between the human and rat occupancy maps. Note the increase across exposure to a maze configuration implying they take increasingly similar routes. **(F)** Average deviation from optimal path (measured in extra maze module visitations) for rats and humans as a function of trial number; humans navigate to the goal with more efficient routes.

### Observations of the behaviour of the RL agents

Examining the trajectories of the RL agents we observed a number of consistent patterns. As expected, model-free agents were relatively unable to adapt to changes in the maze layout; whenever there was a barrier obstructing the learnt route to the goal the model-free agent would remain in a similar region of space and often fail when the new path required travelling away from the goal or around obstacles (Video S3). This behaviour logically follows from the fact that it has no representation of the transition structure and relies on previously cached optimal actions to select which transitions to make. The model-based RL agents generally chose more optimal routes, especially as the trials progressed, although can initially be seen to occasionally make poor choices in paths (Video S4). This is consistent with them requiring an accurate model of the environment in order to conduct a useful tree-search over routes to the goal. However when a change in the transition structure occurred and that model was no longer accurate, they do not have cached values to rely upon and must extensively explore to acquire a model of the new environment they can exploit. SR RL agents initially appear to make similar errors to model-free agents, but adapt more efficiently to the change in the transition structure, for example avoiding deadends after a few trials (Video S5). This is consistent with them updating a stored transition structure using past experience. Thus, unlike the model-based agents, SR agents have a learned set of biases they will fallback to which can aid choice-making after changes in the environment.

### Likelihood analysis of actions reveals rats and humans are both most similar to an SR agent

We next investigated how the human and rat trajectories compared to the RL agents’ representation of optimal actions. To do this, we computed the likelihood of the human and rat behaviour matching each model by restricting the RL agents to follow the biological trajectories. We then used the internal value estimates of the agents to compute a softmax probability distribution over the available actions at each timestep. Using these probabilities to compute the likelihood of the biological data for each agent, we calculated the maximum likelihood parameter estimates for each model’s learning rate and discount factor across individual humans (Table S1) and rats (Table S2).

Comparing the model-free, model-based and SR algorithms, the value representation of the SR agent consistently provided the most likely fit to the biological behaviour (Fig 3A; Likelihood-Ratio (LR) test: SR vs MF for human data ln(LR) = 1911.1; SR vs MB for human data ln(LR) = 538.2; SR vs MF for rat data ln(LR) = 842.0; SR vs MB for rat data ln(LR) = 225.2), with the model-free agent consistently providing the worst fit (MF vs MB for human data ln(LR) = −1372.9; MF vs MB for rat data ln(LR) = −616.9). Consequently, the SR agent was the maximum likelihood model for 70% of the human trials and 60% of the rat trials (Fig 3B). Normalising these likelihoods by trial length and using a uniform random walk as a baseline, we observed this trend was robust throughout the time spent on a maze configuration (Fig 3C) and across individuals (SR vs MF for human data: t(17) = 29.2 *p*<.001; SR vs MB for human data: t(17) = 11.9, *p*<.001; SR vs MF for rat data: t(8) = 13.0, *p*<.001; SR vs MB for rat data: t(8) = 9.6, *p*<.001). We also observed that the agent likelihoods for humans and rats varied across maze configurations (Fig 3D), with a strong correlation between the fits to the biological data (r=.57, *p*<.001).

**Figure 3:**
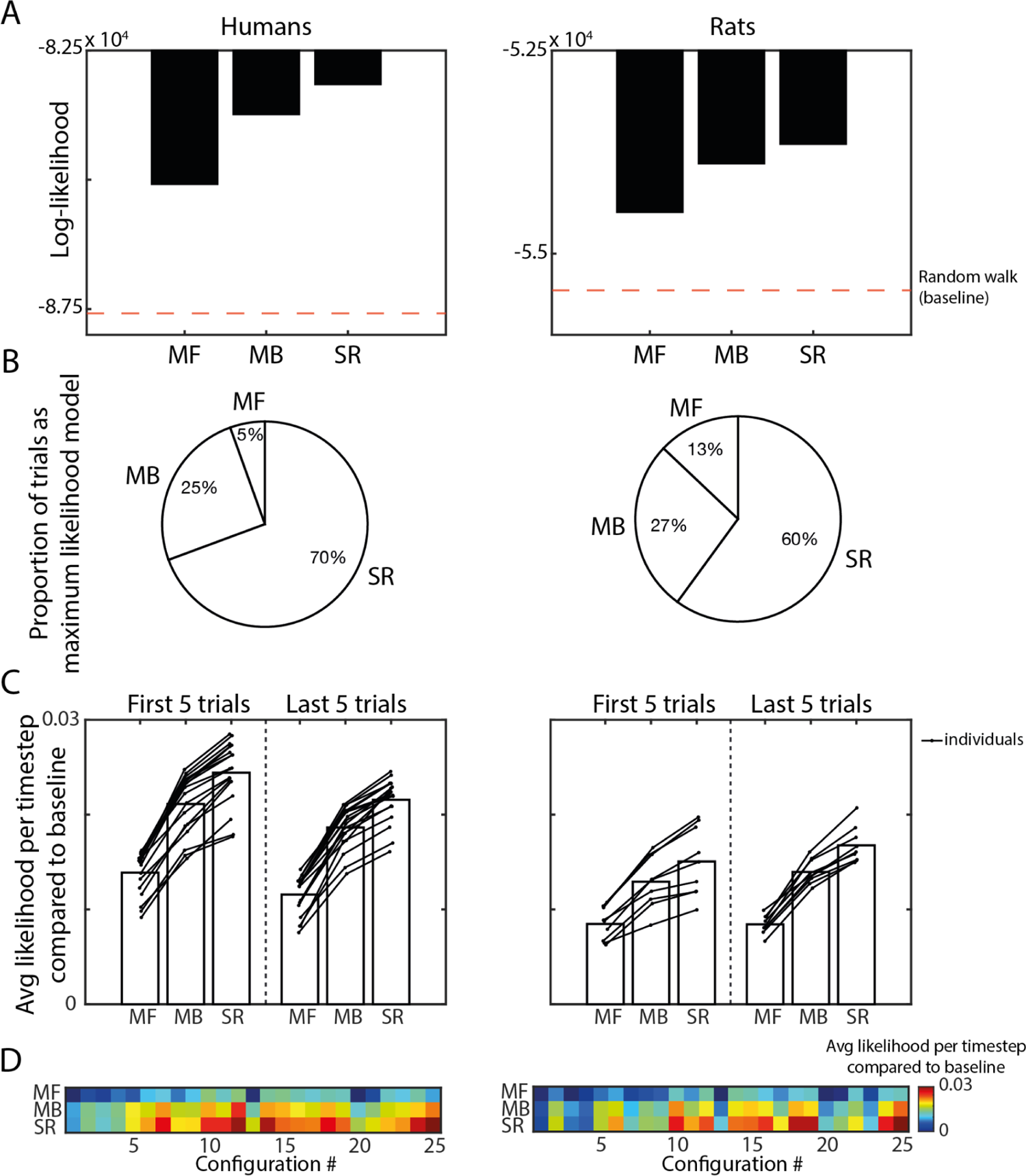
Maximum likelihood analyses of the human and rat trajectories. Likelihood analysis reveals that the behaviour of humans (left) and rats (right) is better predicted by a successor representation agent (SR) than model-based (MB) or model-free (MF). **(A)** The value estimates generated by the SR agent provide a more likely explanation of the biological behaviour than either the MF or MB agents. **(B)** The SR agent was the maximum likelihood model to explain the biological behaviour for the majority of trials. **(C)** This trend is true across all individuals in both species (humans n=18; rats n=9) and robust throughout exposure to a maze configuration. **(D)** Likelihood estimates vary across the maze configurations used, with a strong correlation between the model likelihoods for the human and rat behaviour (r = .57, *p*<.001).

### Simulating agents using parameters derived from the human and rat data reveals closest match to SR agent

To investigate whether these differences in agent likelihoods transfer into measurable differences in the resulting behaviour, we simulated agent trajectories according to each rat and human participant using their individual maximum likelihood parameters (Table S1 & S2). Importantly, these agents were trained on the trajectories taken by that individual on all maze configurations prior to the one being simulated. The agents then carried over all model and value representations learnt across the 10 trials on the simulated maze configuration. To generate the behaviour, the agents followed an ε-greedy policy that linearly decayed from ε=0.1 to ε=0.01 across the 10 trials on a maze configuration. This means for the first trial on a new maze configuration, the agents exploit (i.e. choose the action with maximum expected value) 90% of the time and explore (i.e. choose a random action) on the remaining 10%. Then for each subsequent trial, the agents increase their proportion of time spent exploiting by 1%. To accurately depict the distribution of trajectories generated by an agent under such a policy, we simulated each RL algorithm 100 times per rat and human participant, with the maximum number of state transitions each agent could make set to match the maximum possible for a rat travelling along the grid axes at 20cm/s (i.e. max 45 transitions per trial, 1 transition per second). In the subsequent analyses, individual rats and human participants are compared to the RL agent simulations trained on their individual behaviour, using the maximum likelihood parameters fit to their individual behaviour (Table S1 & S2)

The model-based algorithm generally outperformed the biological behaviour (Fig 4A-B), particularly on the first few trials of a new maze configuration (paired t-test, proportion goal reached on first 5 trials: MB vs humans, t(17)=2.74, p=.014; MB vs rats, t(8)=3.20, p=.013; last 5 trials: MB vs humans, t(17)=0.45, p=.656; MB vs rats, t(8)=2.56, p=.034). The model-based algorithm also consistently outperformed the other RL agents (Fig 4A-B; paired t-test, proportion goal reached, human simulations: MB vs SR, t(17)=22.8, *p*<.001; MB vs MF, t(17)=167, *p*<.001; rat simulations: MB vs SR t(8)=29.0, *p*<.001, MB vs MF t(8)=119, *p*<.001), with the model-free agent performing worst (human parameters: MF vs SR, t(17)=-83.8, *p*<.001; rat parameters: MF vs SR, t(8)=-47.0, *p*<.001). As with the humans and rats, the model-based and SR agents progressively improved throughout the trials on a given maze configuration (Fig 4A-B; first 5 vs last 5 trials, human simulations: MB, t(17)=-40.6, *p*<.001; SR, t(17)=-23.6, *p*<.001; rat simulations: MB t(8)=-18.6, *p*<.001; SR: t(8)=-18.2, *p*<.001). Meanwhile, the model-free agents became progressively worse at reaching the goal (first 5 vs last 5 trials: human parameters, t(17)=35.8, p<0.001; rat parameters, t(8)=16.5, *p*<.001) – indicative of the increasingly complex trajectories required from successive starting positions on a maze configuration. Goal-reaching performance for the RL algorithms varied across maze configurations (Fig C-D), with configurations that had a contradictory optimal policy to the one preceding it seeming particularly difficult (e.g. configurations 4, 8, 13, 21 – see Fig 1C for specific layouts). Conversely, maze configurations that possess a high degree of coherence in optimal policy with the previous configuration (e.g. 2, 7, 25) were consistent with higher levels of agent goal-reaching - due to the improved accuracy of the initial value representations. Ranking maze configuration difficulty by order of goal-reaching performance revealed a significantly more positive correlation between the human and SR agent difficulty rankings, than either the model-based or model-free agents (Fig 4E; paired t-test following Fisher transformation: SR vs MF, t(17)=3.27, p=.004; SR vs MB, t(17)=4.57, p<.001). Similarly, the rat difficulty rankings were significantly more correlated with that of the SR agent than model-free (Fig 4F; (paired t-test following Fisher transformation: SR vs MF, t(8)=2.87, p=.021), with no significant difference to the model-based agent (SR vs. MB, t(8)=1.00, p=.345).

**Figure 4:**
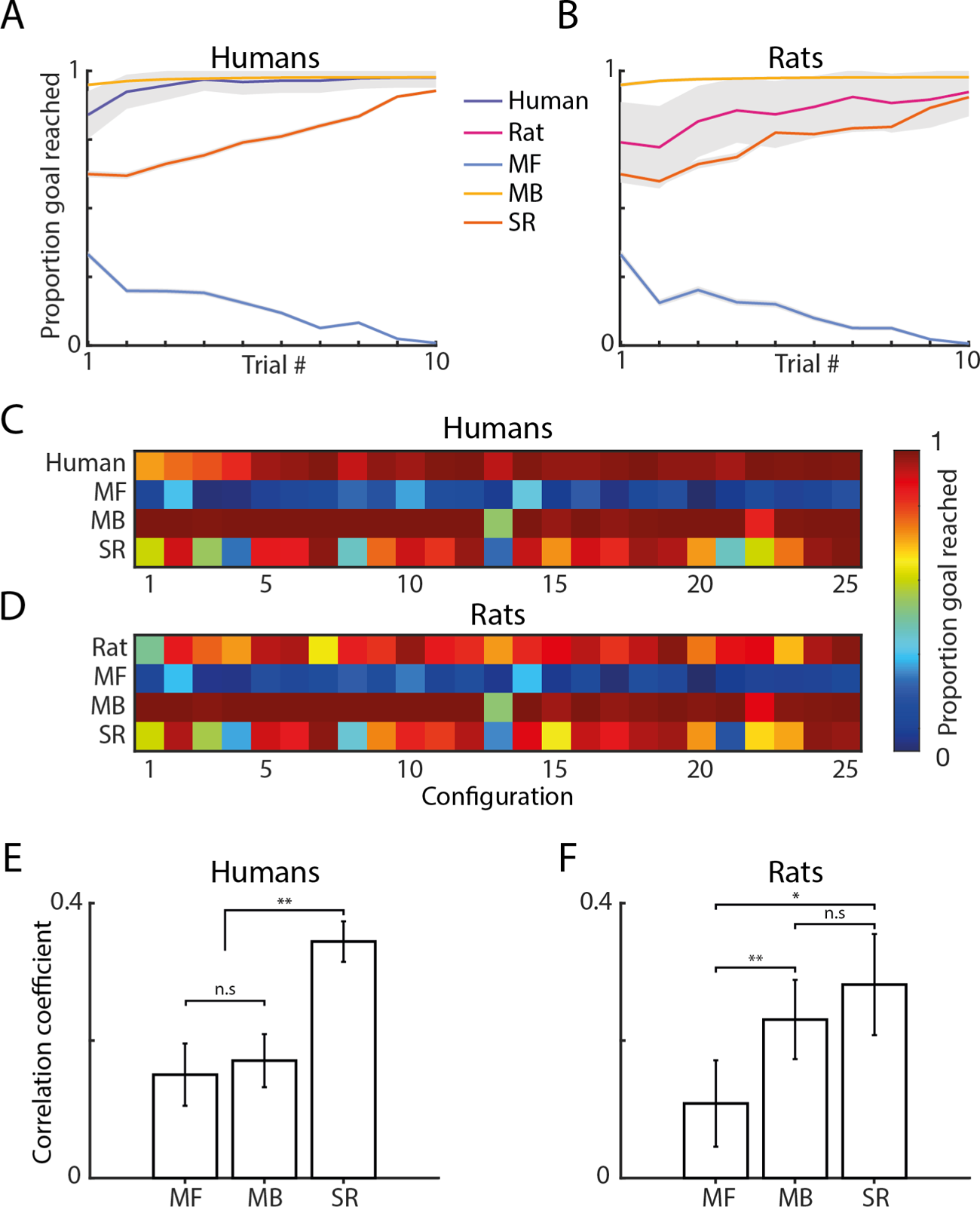
Human and rat performance compared with maximum likelihood reinforcement learning agents. Goal reaching for agents (n=100 per participant / animal) using the maximum likelihood parameters fit to individual human **(A)** and rat **(B)** trajectories. Grey areas indicate standard error from the mean. **(C**-**D**) Goal reaching varied across maze configurations. Using this to rank maze configurations by difficulty reveals a significantly more positive correlation between the human and successor representation (SR) agent’s difficulty rankings **(E)** than the model-based (MB) or model-free (MF) (paired t-test following Fisher transformation: SR vs MF, t(17)=3.27, p=.004; SR vs MB, t(17)=4.57, p<.001; MB vs MF, t(17)=0.35, p=.728). The rat difficulty rankings **(F)** correlated significantly lower with the model-free agent’s than either of the other agent’s (paired t-test following Fisher transformation: MF vs MB, t(8)=-4.33, p=.002; MF vs SR, t(8)=-2.87, p=.021; SR vs. MB, t(8)=1.00, p=.345). Error bars indicate standard error from the mean.

In order to establish whether the routes taken by the rats, humans and RL agents within a maze configuration tended to follow consistent patterns of behaviour, we next quantified each trajectory using diffusivity measures that were inspired by statistical mechanics and the modelling of particles moving in boxes. Specifically, for each trajectory we calculated the linear diffusivity and the sine and cosine of the angular diffusivity (Fig 5A). The linear and angular diffusivities respectively describe the overall directness and direction of the route which vary from trial to trial (Fig 5B-C). Taken together across the entire experiment we see that when the trajectories are quantified this way, unsupervised clustering reveals clear patterns of behaviour for each of the three RL agents (Fig 5D). Given these distinct clusters, we then used the Mahalanobis distance to measure the level of dissimilarity between the biological and agent trajectories per maze configuration. The Mahalanobis distance was used as it accounts for covariance between the diffusivity measures when calculating the dissimilarity. Using these diffusivities to quantify the general shape of the routes taken within a configuration, we found that the trajectories of the SR agents were consistently more similar to the corresponding rat and human behaviour than the other agents (Fig 5E-F; rat simulations: SR vs MF, t(8)=-11.1, *p*<.001; SR vs MB, t(8)=-12.7, *p*<.001; human simulations: SR vs MF, t(17)=-55.7, *p*<.001; SR vs MB, t(17)=-4.21, *p*<.001). Further, the model-free agent was generally the least similar to the biological behaviour across the maze configurations (rat simulations: MB vs MF, t(8)=-2.76, *p*=.025; human simulations: MB vs MF, t(17)=-28.6, *p*<.001; SR vs MB, t(17)=-4.21, *p*<.001).

**Figure 5:**
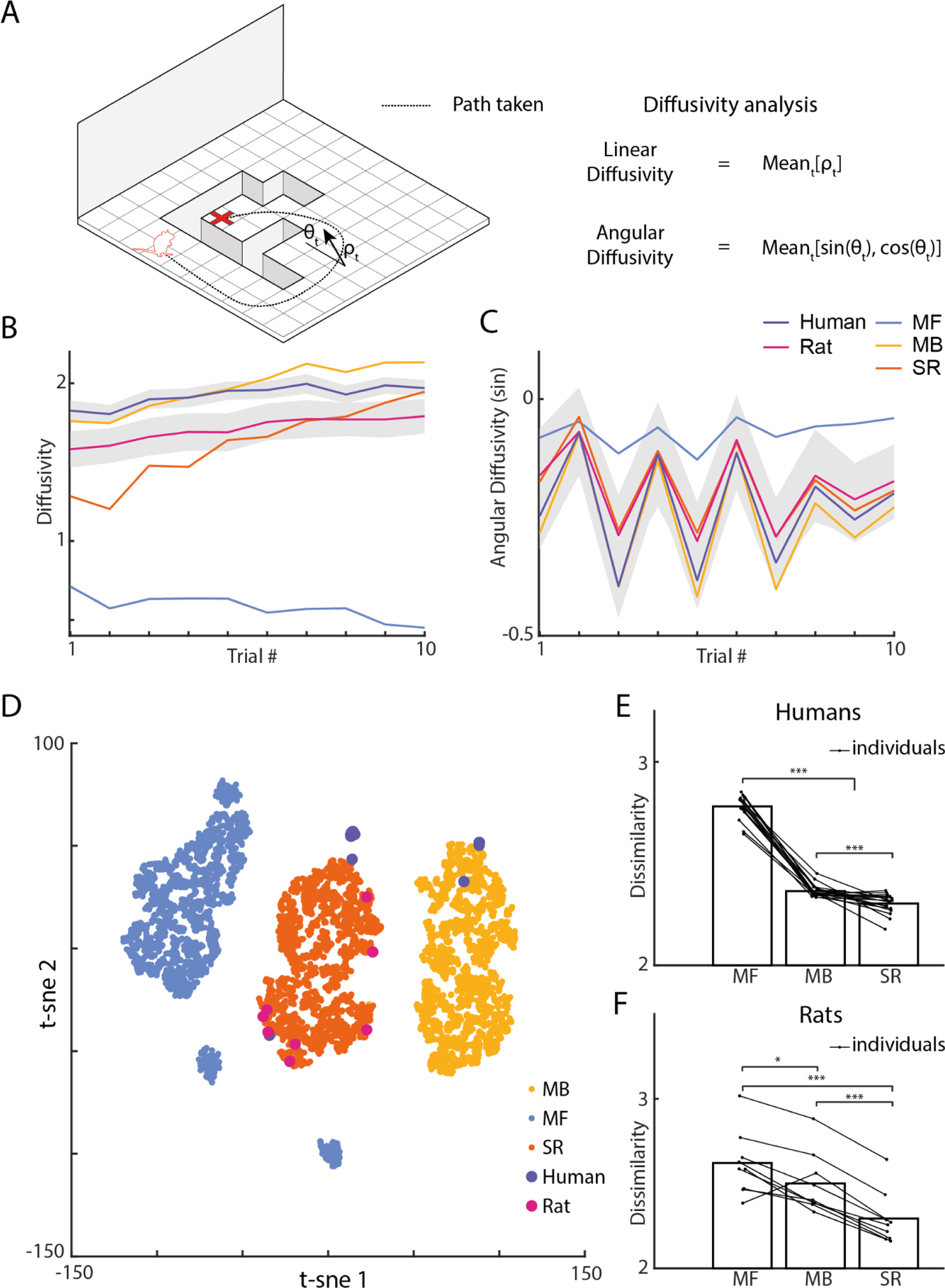
Diffusivity analysis reveals rat and human trajectories are most similar to a successor representation agent. **(A)** Each trajectory was quantified using the average linear diffusivity **(B)** and the sine **(C)** and cosine of the average angular diffusivity. **(D)** Using these metrics to quantify the trajectories on each maze configuration, agent behaviour can be seen to form clusters (shown here via t-sne), where each point represents the average of an individual human, rat or agent over the whole experiment. Note, that for different embeddings a similar pattern emerges. Calculating the Mahalanobis distance between clusters reveals the human **(E)** and rat **(F)** behaviour is more similar to a successor representation (SR) agent than the model-based (MB) or model-free (MF) agents (Humans: SR vs MF, t(17)=-55.7, *p*<.001; SR vs MB, t(17)=-4.21, *p*<.001; Rats: SR vs MF, t(8)=-11.1, *p*<.001; SR vs MB, t(8)=-12.7, *p*<.001).

Finally, to test whether these differences in diffusivity measures across maze configurations directly translated to a physical closeness between individual trajectories, we calculated the minimum path distance between each human/rat trajectory and the simulated trajectories of the agents trained on each individual’s behaviour. Calculating this at every state along a human/rat trajectory and averaging across the length of the trajectory gives a measure of similarity between the biological and agent routes taken. We see that the SR agent trajectories are generally closer to both the human (Fig. 6A; SR vs MB: t(17)=8.32, *p*<.001 SR vs MF: t(17)=28.8, *p*<.001) and rat paths (Fig. 6B; SR vs MB: t(8)=6.44, *p*<.001; SR vs MF: t(17)=26.4, *p*<.001). Interestingly, humans displayed evidence of model-based planning on the early trials of a new maze configuration (Fig. 6C; first 5 trials MB vs SR: t(17)=3.30, *p*=.004; first 5 trials MB vs MF: t(17)=17.0, *p*<.001), with the latter half of trials - when the routes to the goal were longer and more complex - being significantly more SR-like in both humans (Fig. 6C; last 5 trials SR vs MB: t(17)=8.95, *p*<.001; last 5 trials SR vs MF: t(17)=33.6, *p*<.001) and rats (Fig6D; last 5 trials SR vs MB: t(8)=13.4, *p*<.001; last 5 trials SR vs MF: t(8)=24.8, *p*<.001). Viewing how this measure of similarity changes across maze configurations again reveals noticeable variation (Fig 6EF), with a strong correlation in the level agent similarity between the rats and humans (Pearson correlation: ρ = .93, *p*<.001).

**Figure 6:**
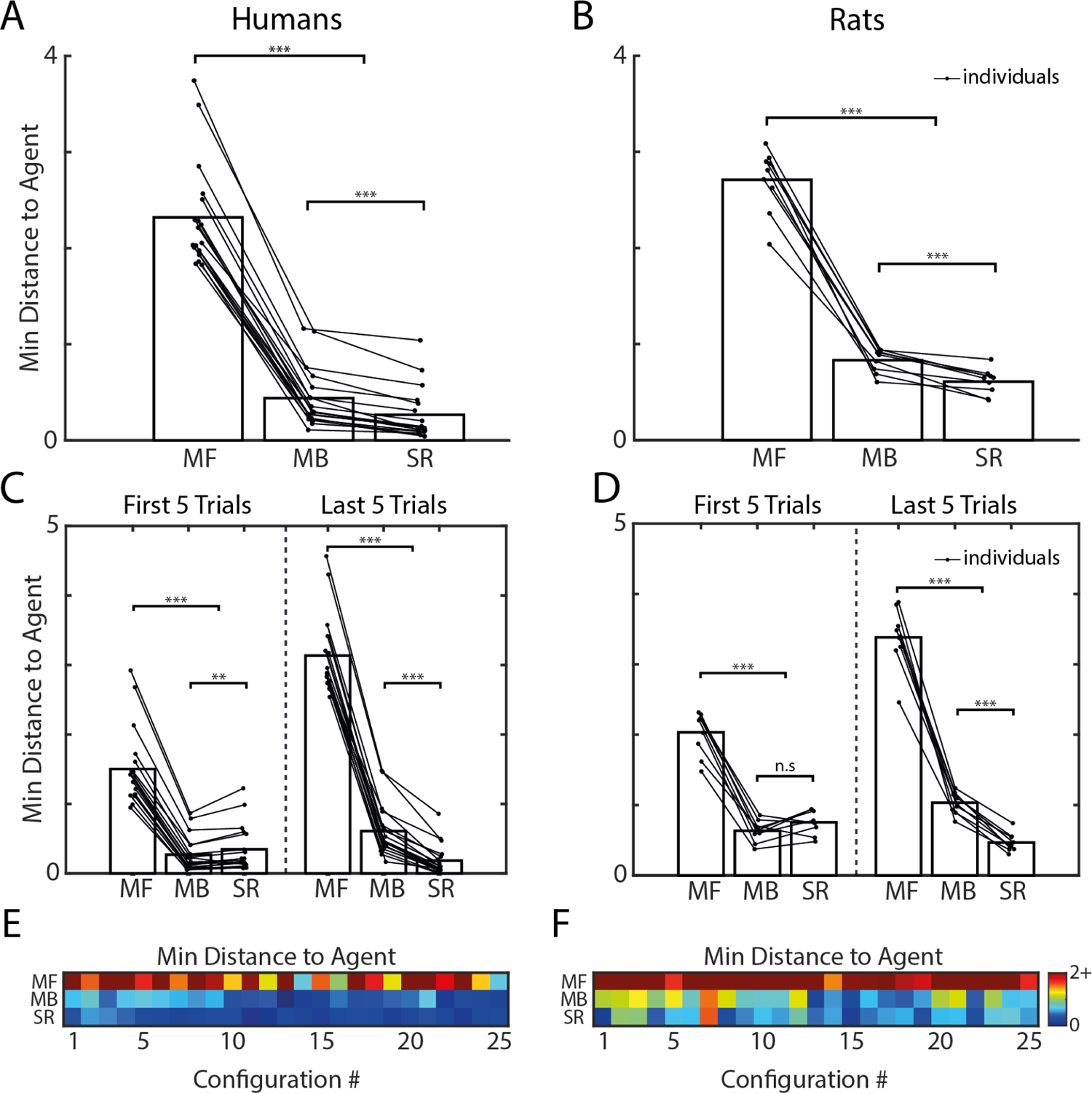
Trajectory similarity analysis identifies SR agent trajectories as closest to human and rat behaviour. Trajectory similarity to the model-free (MF), model-based (MB) and successor representation (SR) agents was measured using the average minimum path distance along each human (left column) and rat (right column) trajectory. **(A)** SR agent trajectories were in general closest to both the human (SR vs MB: t(17)=8.32, *p*<.001 SR vs MF: t(17)=28.8, *p*<.001) and **(B)** rat behaviour (SR vs MB: t(8)=6.44, *p*<.001; SR vs MF: t(17)=26.4, *p*<.001), although **(C)** humans displayed evidence of model-based planning in early trials on a maze new configuration (first 5 trials on a configuration, MB vs SR: t(17)=3.30, *p*=.004; MB vs MF: t(17)=17.0, *p*<.001). Later trials on a maze configuration, which required longer and more complex routes to the goal, were again closest to the SR agent behaviour for both the human (last 5 trials on a configuration, SR vs MB: t(17)=8.95, *p*<.001; SR vs MF: t(17)=33.6, *p*<.001) and **(D)** rat data (last 5 trials on a configuration SR vs MB: t(8)=13.4, *p*<.001; SR vs MF: t(8)=24.8, *p*<.001). The level of similarity to agent trajectories varied across maze configurations **(E**-**F)** with a strong correlation between the humans and rats (Pearson correlation between both matrices: ρ = .93, *p*<.001).

In summary, we used three approaches to compare RL agents to rats and humans: a likelihood analysis of rat and human actions under different agents; the similarity in performance between rats, humans and RL agents trained on the biological behaviour; and the similarity of the resulting trajectories generated by these agents to the individual rats and humans on which they were fit and trained. Our results show that both species match more closely the SR RL agents’ than model-free or model-based agents, with some features of behaviour during early exposure to a new maze configuration being consistent with model-based planning.

## Discussion

To understand the underlying processes that support flexible navigation in rats and humans we compared their navigation performance with three classic instantiations of RL agents in a maze environment using a dynamic layout of barriers. Using a combination of likelihood, performance and trajectory similarity analyses, we find that both rats and humans rapidly adapted to the dynamic environments, producing similar navigation choices and trajectory patterns that most resembled successor representation RL agents. This was most evident for rats, while humans were also found to show some trajectory patterns similar to model-based RL agents in early trials. Our findings provide novel convergent cross-species insights into spatial navigation behaviour and mechanistic understanding of the different choices made when adapting to a changing environment. In doing so we identified a set of metrics that could allow the prediction of future behavioural and neural dynamics across a wide range of methods in humans and animals. We discuss: i) how these results inform our understanding of mammalian navigation, ii) insights into RL models, iii) similarities and differences between rodent and human behaviour, and iv) directions for future research.

### A predictive map for navigation?

Naively, one might view rats as ʻcreatures of habit’, while humans could be considered deep thinkers, mulling over future possibilities. These two stereotypes map, to some degree, onto model-free RL (habit-like) agents and model-based RL (flexible planning) agents. Rather than finding such a dichotomy between rats and humans, we found the behaviour of both species is best captured by an RL agent that creates a predictive map of the environment to guide navigation: the successor representation. The SR has been proposed as an efficient alternative to the relatively inflexible model-free RL and the computationally expensive model-based RL. SR stores a matrix of the possible transitions within the environment and integrates this with information about reward^58^. Recently, it has been proposed that the hippocampus may implement a system similar to a successor representation to create a predictive map to guide navigation^66, 67, 70, 71^. Here we find behavioural evidence to support the proposal that both rats and humans use such a predictive map to guide flexible navigation behaviour. This match of the rodent behaviour to SR agents is consistent with evidence that rats can carefully evaluate different options for navigation^72^.

A range of previous experiments comparing human behaviour to RL agents have generally focused on a competition of model-free vs model-based agents to capture behaviours in small conceptual state spaces with 2-step transitions. These have found evidence for model-based planning in humans^68, 73, 74^. Using a much larger state space with potential for recursive transitions (i.e. loops leading back to the same state), we extend this approach into a more complex and naturalistic framework. Our findings add to recent evidence that the choices of humans are best explained by a combination of SR and model-based behaviours^63–65^. Since the SR encodes the environment’s transition structure, it is itself a transition model that can be leveraged for intuitive planning^75^ or more explicit planning procedures such as a tree search - which may partially explain trajectories observed during hippocampal replay^23, 44, 76–79^. Given that the model-based learner will generally improve the accuracy of its learnt model as it has more experience of a maze layout, it might be surprising that we observed a greater match to SR agents during the second half of trials on a new maze configuration. However, the model-based planning mechanism of simulating possible future paths is considerably more resource intensive than drawing upon a cached knowledge of past behaviours gleaned from experience. Thus, for a metabolically constrained learning system it would be more efficient to fallback on simpler processing mechanisms when they reach a certain threshold in terms of performance (e.g. maximising reward and/or minimising uncertainty in expected reward) - which is supported by evidence of in both rodents and humans^50, 80^.

A few studies have explored human navigation in VR environments and compared navigational choices with RL agents, reporting that behaviour matches a mix of model-based and model-free choices^45, 49^. For example, when paths are short but decision times are longer, model-based RL agents were found to better match human behaviour^49^. However, these past studies did not compare performance with a successor representation, nor were the trajectories examined in relation metrics such as the diffusivity and physical closeness to understand the match to different RL agents. Here we show that navigation in humans is most similar to SR and that using trajectory information is useful in providing convergent evidence to understand this.

In our experiment the model-free RL agents showed poor adaptation to the changes in maze layout. Rather than improving over trials, performance declined. This could be accounted for by our task structure; the minimum trajectory length increased as trials progressed, requiring longer and more complex routes to the goal. Our maze configurations were designed to be simple but to include dead-end zones and/ or regions where the barriers extended around the goal zone requiring extended trajectories away from the goal to then reach it. It is possible with different layouts model-free learners would succeed more efficiently. Understanding how the structure of the task and state space leads to the emergence of different policies is an important question for future research.

A key observation in our data is that it is not sufficient to conclude on the basis of overall performance which simulated agents will best fit the biological agent’s data. While the model-based RL agent performed best and was closest overall to the human performance, the SR RL model produced the greatest match in terms of the proportion of trials as the maximum likelihood model. This is because the patterns of choices made by the model-based RL fail to capture aspects of choice and trajectory patterns in the rats and humans to the same degree as the successor representation agents. This highlights the utility of an environment in which a wide diversity of trajectories can be achieved by the rats and humans to allow models to be discriminated.

### Similarities and differences in the flexible behaviour of rats and humans

Past research has suggested that rats do not always optimally adapt to selecting appropriate alternative routes when navigating^9^ and can take time to adjust to such changes to select the optimal route^14, 81^. Similarly, humans can struggle to take optimal shortcuts when presented with options^8, 19, 20^. Here rats and humans had to reach the correct goal from a set of 100 possible locations within a 45 second time limit. Since the maze transition structure was re-configured every 10 trials, achieving this was non-trivial. Despite this, both species were able to reach the goal on the first trial of a new layout on many trials. This parallels recent evidence from mice learning new paths remarkably fast in a large labyrinth^82^. In several cases we saw examples of routes near the optimal path on the first attempt for both species (e.g. see Videos S1&2). Moreover, the occupancy correlation between rats and humans was relatively high even from the first trial, and improved as performance increased across trials in a configuration. These results show that our Tartarus maze, with its visual access to landmarks, boundary geometry and canyon-styled barriers, provides a useful assay for goal-directed navigation across two species, revealing a remarkable similarity in the patterns of navigation across species.

Despite similarities in behaviour, there were noticeable differences between species. Humans were more successful at adapting to the changes in maze layout and learning, while rats spent more time on the perimeter of the maze’s edge. The overall difference in learning likely relates to the physical differences in our maze used between rats and humans (real vs VR^83^) and the biological differences between species (e.g. differences in vision, movement, whisking, olfaction, grooming and predator/prey status). The current models assume optimal routes minimise distance. However, rats will also need to avoid predators, thus selecting certain routes that are safer may also drive route choice^14^. Rats also need to find suitable and safe places to groom their fur. These factors may underlie the generally poorer fit of the models to rats than humans. Additionally, while fog was used in the human VR to better match the visual acuity and depth perception between rodents and humans^119^, the rats had visual access to more of the maze during the experiment - with recent evidence in humans suggesting this can bias strategies towards a SR^52^. Further research would be needed to disentangle the various contributions that give rise to the differences we observed.

### Benefits of a dynamic open field environment with barriers

Prior studies examining navigation in mazes have generally either used track-based or open field environments^28, 84, 85^. While open field environments place more demands on self-localisation and vector-based navigation^78, 86, 87^, mazes with tracks enable testing the effect of blocked paths and shortcut behaviours^6, 9, 81, 84^. By contrast the Tartarus maze places demands on both vector based navigation and the capacity to take detours and shortcuts, as occurs with much of the terrain in the real world. The recent development of the honeycomb maze for rats^88^ provides a parallel approach to self-localisation and obstructed paths to goals, where rats sequentially navigate to a goal over a number of hexagonal platforms that are made available in pairs until the goal platform is reached. Such an approach allows for a precise assessment of choice options at different time points, whilst placing demands on self-localisation in relation to distal cues. While the tartarus maze also demands choices and navigation to distal cues it allows continual, often ballistic trajectories to be taken to the goal, mimicking naturalistic behaviours that enable integration with more ethological approaches to navigation^89^.

A number of rodent studies have examined how maze layout and changes in layout relates to exploration behaviour^82, 90–94^ or escape behaviour^14, 95^. Here we found rats and humans rapidly adapted to changes in the maze structure, by exploring the new layout. By matching to RL models it is possible to provide a more mechanistic account of how goal-directed behaviour is organised during the navigation of a dynamic environment. In the case of Rosenberg et al. (2021)^82^ mice had to learn the paths in a maze with a large number of options. Akin to our task, learning was rapid. This differs from many non-spatial learning tasks where learning is typically slow (see Rosenberg et al., 2021). Other recent rodent studies exploring navigation behaviour have shown the capacity to model behaviour in goal learning and homing vectors for safety^14, 87, 94, 96^. Such studies highlight the value in modelling to understand the mechanisms guiding behaviour. Here, we demonstrate the added benefit of modelling behaviour with simulated agents, examining the trajectory properties (e.g. diffusivity) and comparing across two different species. Across many studies distal cues are kept constant allowing for rapid learning of the new layout. Manipulating these distal cues would be an interesting direction for future research.

Prior human virtual navigation studies exploring flexible navigation behaviour have tended to involve complicated VR environments that would likely be too demanding for rodents to learn^10, 12, 13, 20, 45, 52, 97, 98^. Here, we sought to recreate an environment that would challenge human participants generating sufficient variation in performance and to allow comparison to rodents within the same maze structure. Being able to integrate behavioural data from humans, rodents and RL agents, opens the possibilities for incorporating data from a wide array of neuroscience methods in humans and rodents. Recent students have shown the utility of this approach^29–31^. A recent study by Zhu et al. (2021)^52^ highlights the benefit of examining eye-movement dynamics during the navigation of virtual environments, using a similar head-mounted display to our human VR, but where the whole transition structure was visible to look at. Their results show patterns of eye-movements that sweep across key points in mazes, maximally important for planning, showing forward sweeps to the goal, as well as backward sweeps from the goal. Furthermore, they show evidence that patterns in eye-movements that scan relevant available transitions relate to SR agent performance. Future work with eye-tracking integrated into our human task would be useful to study the selection of sub-goals and eye-movements after changing the maze layout; could eye-movements predict future choices of route and the match to different RL agents in subsequent behaviour? Eye-tracking in rodents is a bigger challenge but may also hold some promise^99^. Finally, it may also be useful to explore the search behaviour of RL agents, humans and rats in relation to models of utility and biased search^100^. Such explorations would be interesting to examine in mazes ranging in complexity, visibility (fog-levels) and frequency of re-configuration. Based on our results we would predict that human behaviour would match model-based agents more in rapidly changing environments and environments where they can see more of a complex layout that would benefit from deliberating over the options.

### How might the RL agents be improved?

The learning efficiency of RL agents could be improved using offline replay of randomly sampled past experiences^44, 60, 64, 101^. These replays are typically implemented between agent timesteps, and the manner in which they are sampled can further accelerate learning by prioritising the most useful learning experiences to replay^76^. Prioritised replay also has strong parallels to the phenomenon of hippocampal replay of place cell activity during sleep or quiescence^78, 102, 103^. However, in this study we did not implement agent replay in order to keep the value representations, and consequently the likelihoods of agents, deterministic. An alternative way to improve the goal-reaching of agents could be through improving their exploration policy. The agents simulated here relied on an ε-greedy policy through which exploration is driven purely by chance. However, methods that include curiosity^104^ or uncertainty in the value function^105, 106^ could be used to guide more efficient exploration of a new maze configuration and consequently lead to faster learning. Finally, navigation using an options-framework might allow for more efficient navigation^107^; rather than planning step by step, efficient navigation in our maze configurations can be achieved by selecting a clockwise vs a counter-clockwise path to the goal. Being able to exploit a hierarchical segmentation of the environment might allow RL agents to better approximate human and rat behaviour (see e.g. Balaguer et al., 2016). Further, the points where agents switch between different options may be able to predict where rats and humans would pause in the maze^72^. More broadly, new approaches to RL^109^ and deep learning methods may provide new ways to examine navigation^40, 110^, as well as integrating our approach with biologically-inspired network models that seek to explain neural dynamics during navigation^111, 112^.

### Exploring the neural substrates of a predictive map

Recent neuroimaging in humans has shown that activity in hippocampal and connected regions tracks the modelled parameters from a successor representation^62, 63, 65, 67, 77^. Convergent evidence in rodents suggests that the place cell activity in the dorsal CA1 of rodents may operate as a successor representation^66, 67, 70, 113, 114^. Our protocol would allow for evidence from both rodent and human data to be integrated within a single framework to consider how patterns in the data may interrelate across species and in relation the parameters from RL modelled agents. Evidence from other recent approaches shows the utility of such an approach^29–31^. Our recent analysis of CA1 place cell activity found little evidence for changes in the place field maps when the state space changed due to blocked doorways in a 4-room maze^109^. However, it appears the changes in layout are evident during hippocampal replay events, where paths activated follow the new layout, perhaps consistent with MB-RL search patterns^79^. One consideration for future research will be to explore changes in neural activity linked to particular strategies that might occur over trials or even within a route. For example, one might predict a shift to a more striatally mediated strategy, linked to model-free RL, if the number of trials for a configuration was increased^50, 56^. Shifts between the engagement of different structures to guide control may also occur alongside shifts within structures^50, 80^. For example, the hippocampus might be involved in simulating paths via replay to guide behaviour in highly dynamic environments, but shift to a more cached expression of the stored hippocampal map once the possible paths have been repeatedly experienced.

Recent work modelling multi-scale SR agents^116^ has shown patterns similar to the goal distance tuned activity of CA1 cells in navigating bats^117^. Might such patterns emerge in our task? An important step in better understanding the neural systems for navigation would be to examine the impact of temporally targeted inactivation of the hippocampal regions, as well as the prefrontal cortex which is thought to support route planning^37^. Such an approach would provide more causal evidence for the role of brain structures for supporting our task. More broadly, the task we have developed could be adapted for study with a range of other species which have been examined in isolation: ants, bees, drosophila, bats, birds and other primates. Integrating across invertebrates and vertebrate species may further our understanding of the common mechanisms for goal-directed behaviour and adaptations that occurred through evolution.

## Conclusion

In summary, we found that rats and humans both display behaviour most similar to a successor representation RL agent, with humans also showing some behaviour matching model-based planning. Future work exploring single-unit recording or disruption to neural activity may be useful in revealing how distance to the goal may be coded, as past studies have failed to dissociate path and Euclidean distance. Moreover it will be useful to examine how neural activity in humans and rodents relates to the parameters from RL agents with behaviour adjusted to match the humans and rats. More broadly, the approach provided here could be adapted to compare behaviour across a range of species and different RL models to help understand the broad spectrum of navigation behaviours shown by the diverse species on our planet.

## Acknowledgments

We would like to thank Célia Lacaux and Charles Middleton for their help building the maze. WdC was funded by an EPSRC CASE studentship with Deepmind Ltd. NN and HJS received support from the European Union’s Horizon 2020 Framework Programme for Research under Marie Sklodowska-Curie ITN (EU-M-GATE 765549). CB is funded by Wellcome SRF (212281/Z/18/Z). HJS received funding from Deepmind Ltd.

## Author contributions

Conceptualisation, WdC & HJS; Methodology, WdC, FZ, SR, DB, RG, ED, CB & HJS; Formal Analysis, WdC; Investigation, WdC, NN, EMG, CG, JL, LF and CN; Writing - Original Draft, WdC & HJS; Writing - Review & Editing, WdC, NN, CN, DB, RG, ED, CB & HJS; Visualisation, WdC, FZ, HJS; Supervision, CB & HJS.

## STAR Methods

### RESOURCE AVAILABILITY

#### Lead contact

Further information and requests for resources should be directed to and will be fulfilled by the Lead Contact William de Cothi

### Materials availability

This study did not generate any unique reagents.

### Data and code availability

All data and code have been deposited at zenodo and are publicly available as of the date of publication. DOIs are listed in the key resources table.

### EXPERIMENTAL MODEL AND SUBJECT DETAILS

Nine adult male Lister Hooded rats were handled daily (at start of training: 10-20 weeks old, 350-400 g) and housed communally in groups of three. All rats were subjected to a reverse light-dark cycle (11:11 light:dark, with 1 hour x2 simulated dawn/dusk) and were on food-restriction sufficient to maintain 90% of free-feeding weight, with ad libitum access to water. The free-feeding weight was continuously adjusted according to a calculated growth curve for Lister Hooded Rats^118^. Six rats were naive, while three rats had previously been trained for 2-3 weeks in a shortcut navigation task for a different maze setup. The procedures were conducted according to UCL ethical guidelines and licensed by the UK Home Office subject to the restrictions and provisions contained in the Animals Scientific Procedures Act of 1986.

For the human version of the task, 18 healthy participants (9 female; aged = 24.6 ± 5.9, mean ± sd) were recruited from the UCL Psychology Subject Pool and trained to navigate to an unmarked goal in a virtual arena of approximately the same relative proportion as for the rats. All participants gave written consent to participate in the study in accordance with the UCL Research Ethics Committee.

## METHOD DETAILS

### General methods

Navigation was tested in a large square environment with a fixed hidden goal location and a prominent directional black wall cue in one direction (Fig. 1, Video S1 and S2). The maze was divided in a 10×10 grid of moveable sections that could either be removed, leaving impassable gaps to force detour taking, or added, creating shortcuts. During training, all maze modules were present. Rats, humans and RL agents were trained to reach the goal within a 45s time limit, (Fig 1A). During the testing phase of the experiment, maze modules were removed to block the direct route to the goal (Fig 1B). Humans (n=18), rats (n=9) and agents were tested on the same sequence of 25 maze configurations each with 10 trials in which a set of defined starting locations were selected to optimally probe navigation (Fig 1C). These maze configurations were generated from a pilot testing with 9 rats and the configuration sequence chosen maximised the differences in the layouts between trials. The starting positions on each maze configuration gradually increased in the required tortuosity (path distance / Euclidean distance) of the shortest path to the goal to test complex trajectories whilst keeping the rodents motivated.

Upon reaching the goal module, rats and humans had to wait 5s to receive their reward. Human participants were rewarded with a financial bonus and rats received chocolate milk delivered in a well (supp. fig 1). In order to better match the visual acuity and depth perception between rodents and humans^119^, a thick virtual fog lined the floor of the maze enabling them to only see adjacent maze modules and the distal black wall cue (supp. fig 2; Video S2). Modules were made visually indistinct to avoid humans counting them when traversing the space. Human participants were informed that reward was hidden in the environment and that their task was to maximise their financial return as quickly and efficiently as possible. The human and rat trajectories were discretised into the underlying 10×10 modular grid (Fig 2A-B) in order to facilitate comparison between each other and the RL agents.

In all versions of the experiment, the environment (raised off the floor) consisted of a 10×10 grid of maze modules. These modules could be removed from the grid in order to form impassable barriers in the environment. One of the modules was rewarded and thus was the location of the goal in the maze. Navigation was facilitated by a single distal cue consisting of a black curtain that spanned the majority of one side of the maze. The goal was kept in the same position with respect to this distal cue throughout all versions of the task. All participants, rats and learning agents were initially trained to navigate to the goal module on the open maze, without any maze modules removed. Once trained, they were all put through the same sequence of 25 maze configurations, with the same sequence of starting locations on each configuration.

### Rodent methods

All procedures were conducted during the animals’ dark period. The experiment was carried out in a custom-made modular 2×2m square maze composed of 100 identical square platform tiles elevated 50cm above the ground (supp. fig 1). The maze was constructed from Medium Density Fibrewood, with the platforms painted in grey. Each platform contained a plastic well (32mm diameter) at its centre, which could be attached to a polymeric tubing system installed beneath the maze. This tubing allowed the experimenter to reward the rat at the goal module filling the well with chocolate milk (0.1 ml). Importantly, all modules in the rodent maze were identical in appearance and construction with chocolate milk rubbed into the well of non-goal modules to lower reliance on olfactory navigational cues. The maze was surrounded on all sides by a white curtain, with a black sheet overlaid on one side to provide a single extra-maze cue. To ensure that no other cue could be used by the animal (uncontrolled room cues, olfactory traces on the maze) the black sheet was rotated 90° clockwise between sessions. The goal module was always in the same position with respect to this cue. Moreover, the experimenter stayed next to the maze inside the curtained area throughout all sessions, his positions relative to the goal were randomised.

### Familiarisation

During the first day, the rats received a small amount (0.1ml per rat) of chocolate milk in the home cage to decrease neophobia in the maze. For the subsequent two days, each rat underwent two 15 minute maze familiarisation sessions, in which the rat was placed at the centre of the maze and would forage for pieces of chocolate cereal (Weetos) scattered throughout the maze. More cereal was concentrated in the centre to encourage the animal to be comfortable in the middle of the maze.

### Training

Training consisted of two stages, rats were given 2 training sessions per day. In each training trial the rat had 45s to find the goal module.

For stage 1 of training the goal well was filled with 0.1ml of chocolate milk and the rats were initially placed on the modules adjacent to the goal, facing the goal. If the rat made two consecutive direct runs to the goal (without exploration of other parts of the maze), the next trial began one module further away from the goal. Conversely, if the rat failed two consecutive training trials, the next trial began one module closer to the goal until the rat was back at the goal-adjacent modules. On day 1, this procedure was continued until 15 min had elapsed.On the following days, the number of trials was fixed to 16. This procedure was followed every day until the rat was able to make direct runs from the far edges of the maze.

Stage 2 was similar to stage1 but a delay in the release of chocolate milk was introduced. This delay started at 1s and was gradually increased until the rat could wait at the goal location for 5s before the chocolate milk was released. Furthermore, the rat’s starting position and orientation were randomised. The number of daily trials could be increased up to 25. This procedure was followed until the rats were able to successfully navigate directly to the goal and on at least 90% of trials. The training phase took on average 24 sessions.

### Tartarus Maze testing

Rats were run on the 25 maze configurations. For each maze configuration, rats were given 10 trials where they were placed by hand at the starting positions indicated in figure 1. Trials were 45s long and rats were required to navigate to the goal within this time and wait for 5s in order to receive the reward (0.1ml of chocolate milk). If the rat failed to reach the goal, they received no reward and were placed by hand at the next starting location. The rats would usually complete 3 configurations per day. At the beginning of each day, rats were given a brief reminder session that consisted of 5 trials from phase 2 of the training phase.

### Human methods

Participants were reimbursed for their time as well as a bonus of up to £25 for good performance in the testing phase. Participants experienced the virtual environment via a HTC Vive virtual reality headset whilst sat on a swivel chair. They were able to adjust movement speed using the HTC Vive controller and movement direction was controlled by the participant’s orientation on the chair. Upon successful navigation to the goal module, participants were informed of their financial reward along with the presence of a revolving gold star (supp. fig 2) at the goal location. In accordance with the rodent experiment, navigation was aided by the presence of a black distal cue that took up the majority of one of the walls. Goal location, maze configurations and starting positions were all defined with respect to this distal cue and were identical to the rodent experiment. Importantly, a fog lined the floor (supp. fig 2, Video S2) of the maze to prevent the participants from understanding what maze modules were missing until they were at adjacent locations. This also provided a better match to visual information available to the rats - which are known to have less visual acuity and binocular depth perception ^119^. Seamless textures were applied to the floor and walls of the virtual environment, and these were rotated every 10 trials to prevent them from being used as extraneous cues for navigation.

The experiment took place over four sessions on four consecutive days. The majority of the first session was usually spent training the participants to navigate to the goal module. To accelerate this learning process, the participants were initially able to see a revolving gold star in the goal location. As they progressed through the training session the star became increasingly transparent until invisible, with the star only appearing again upon successful navigation to the goal module. Along with the decreasing visibility of the goal, the participants’ starting positions were moved progressively further from the goal in a similar manner to the rat training phase. All training and testing trials were 45s in length. Training was terminated when the participants were able to navigate to the hidden goal on at least 80% of trials after being randomly placed at the far edges of the environment. Mean time to complete this training was 41 ± 21 minutes. In order to make the participants’ experience similar to that of the rodents, they were not given any explicit information about the nature of the task - only that financial reward was hidden in the environment in the form of a gold star and their task was to maximise their financial return as quickly and efficiently as possible.

The testing took place over the remaining sessions and on average lasted 125 ± 25 minutes, with participants encouraged to take short breaks every 10-20 trials to reduce virtual reality sickness. At the beginning of each testing session, participants completed a short reminder task, which consisted of 5 trials from the end of the training phase.

### Reinforcement learner simulations

Reinforcement learning seeks to address how an agent should choose actions in order to maximise its expected accumulated reward yielded from future states, which is known as the value function *V*:

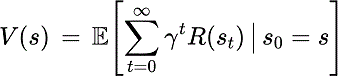

The parameter γ is a discount factor that determines the timescale of how motivating future rewards are, such that for γ < 1 the agent exponentially discounts future rewards^53^.

The reinforcement learning agents were implemented in a 10×10 grid world environment, with each state in the grid world corresponding to a maze module in the human/rat versions of the task (see Fig. S1&2). Thus, unlike the humans and rats, the agents were not explicitly required to self-localise with respect to distal cues, rather they were given absolute knowledge of their current location (state) on the maze in the form of a one-hot vector (a vector with a ʻ1’ in the element corresponding to the current state, with all other elements in the vector being ʻ0’). Upon receiving this information pertaining to its current location, the agent was able to choose actions (i.e. up, down, left, right) which transition it to adjacent states, with the ultimate aim being to choose a sequence of states leading to the goal. Crucially, the way in which an agent chooses this sequence of states is different for the model-free, model-based and successor representation algorithms - which are explained in more detail below. At the beginning of the experiment, all agents were endowed with the optimal policy on the open maze to simulate the training phase undertaken by rats and humans. They were then run consecutively on the 25 maze configurations, using the maximum likelihood parameters fit to each individual rat or human participant’s data. For a given individual rat or human, agent behaviour was simulated on each maze configuration by first training the agent on all of that individual’s trajectories (in the same sequential order) prior to the configuration being simulated. Agents then carried over all models/value representations learnt during their 10 trials on the maze configuration being simulated. Hence, the simulated behaviour of agents was never trained using the human/rat trajectories on the configuration being simulated, only the trajectories on all configurations prior. Each type of agent (model-free, model-based and successor representation) was simulated N=100 times per rat/human, using an ε-greedy policy with ε linearly decaying from ε = 0.1 to ε = 0.01 across the 10 trials on a maze configuration. This means that on a new configuration the agents initially chose the greedy action 90% of the time and a random action the remaining 10% of the time (in order to manage the exploration-exploitation tradeoff), with the agents increasing the proportion of greedy actions they take by 1% on each subsequent trial. Due to the behavioural variance introduced by this policy, each algorithm was implemented 100 times for each rat/human to produce the distribution of behaviour used for the comparison with biology. In the subsequent analyses, each individual rat or human was compared to the simulated agents trained on their behaviour, using the maximum likelihood parameters fit to their behaviour.

### Model-free agent

The model-free method uses the state-action value function instead of the state value function *V*.

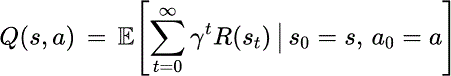

State-action values were learned using the Q-learning algorithm ^55^ combined with an eligibility trace^53^. The eligibility trace is a decaying trace of recently taken state-action pairs. Specifically, after taking action *a_t_* in state *S_t_* and transitioning to state *S_t+1_* where it receives reward *r_t_*, the agent will first decay its eligibility trace *e* - a matrix with the same dimensions as *Q*:

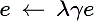

where λ = 0.5 is the eligibility trace decay parameter and γ is the discount factor of the value function in tables S1 & S2. Next, the model-free agent will update its eligibility trace:

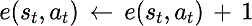

before finally updating the state-action values according to:

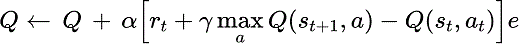

where α is the learning rate in tables S1 & S2. Under a greedy policy, the model-free agent at decision time will choose the action *a* in state *S* with the highest state-action value *Q (s, a)*. If multiple actions with the same maximal value exist, then the agent samples from these with equal probability. The eligibility trace *e* is set to zero at the beginning of each trial

### Model-based agent

The model-based agent is provided with an internal 10×10 binary grid representation of which maze modules are present or not in the environment. Every state in the agent’s model *x* corresponds a module in the maze (see Fig 1A-B); as it transitions through the environment, it updates the internal model at every timestep according to the adjacent states *s*’.

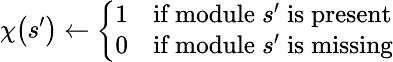

At decision time, the model-based agent uses its model and to plan the shortest route to the goal from each possible next state. Shortest routes were calculated using an A* tree search algorithm ^120^. In the event of multiple equally short routes to the goal, their respective actions were sampled with equal probability.

### Successor representation agent

The SR somewhat combines parts of model-free and model-based learning^59, 60^ by using experience to learn a predictive map *M* between the states in an environment. For a one-step state transition matrix *T*, the predictive map is equivalent to the discounted sum of future state transitions:

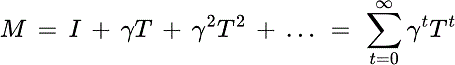

This discounting of transitions means *M* can be readily combined with a separately learned reward *R* associated with each state *S* in order to explicitly compute value.

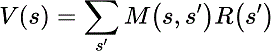

The SR agent uses temporal-difference learning and eligibility traces to update the successor matrix *M*^121^. After transitioning from state *S_t_* and to state *S_t+1_*, the agent will first decay its eligibility trace *e* - a vector with length equal to the number of states in the environment:

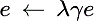

where λ = 0.5 is the eligibility trace decay parameter and γ is the discount factor of the value function in tables S1 & S2. Next, the successor representation agent will update its eligibility trace:

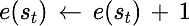

before finally updating the successor representation ^121^:

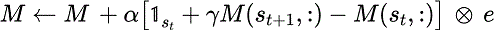

where ⊕ indicates an outer product. This can then be combined with the state-rewards *R* at decision time to compute the value of prospective future states.

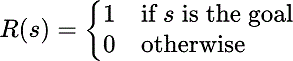

Under a greedy policy, the successor representation agent at decision time will choose the next available state with the highest value. If multiple available states exist with equatlly high values, then the agent samples from these with equal probability. The eligibility trace *e* is set to zero at the beginning of each trial.

## QUANTIFICATION AND STATISTICAL ANALYSES

Optimal paths were calculated using the A* tree search algorithm^120^ in the 10×10 grid state space, with path length measured in terms of state visitations. Occupancy correlations were calculated using the Pearson correlation between the proportion of time spent in each state of the 10×10 grid state space. One- and two-sample t-tests were implemented using MATLAB’s ttest and ttest2 functions.

Likelihoods were calculated by inputting individual human/rat state trajectories to the RL agents and calculating the internal value estimates of the available state transitions conditional on the human/rat’s past trajectories. These value estimates were used in a softmax function to calculate at each time point, the probability that the agent would take each of the available actions conditioned on the human/rat’s past. Maximum likelihood parameters were estimated using MATLAB’s fmincon function to minimise the negative log-likelihood.

Mahalanobis distances were calculated using MATLAB’s pdist2 function on the diffusivity metrics for the humans, rats, model-free, model-based and successor representation agents.

The minimum path distance analysis used an individual human/rat trajectory as a reference trajectory. At each time point along that trajectory, the A* tree search algorithm^120^ was used to find the shortest path distance on the maze configuration to the agent trajectories trained from that individual human/rat’s behaviour. Averaging along the length of the trajectory then gives a measure of similarity between that reference trajectory and the simulated agents.

## Supplementary Figures, Tables and Videos

**Supplementary Figure 1:**
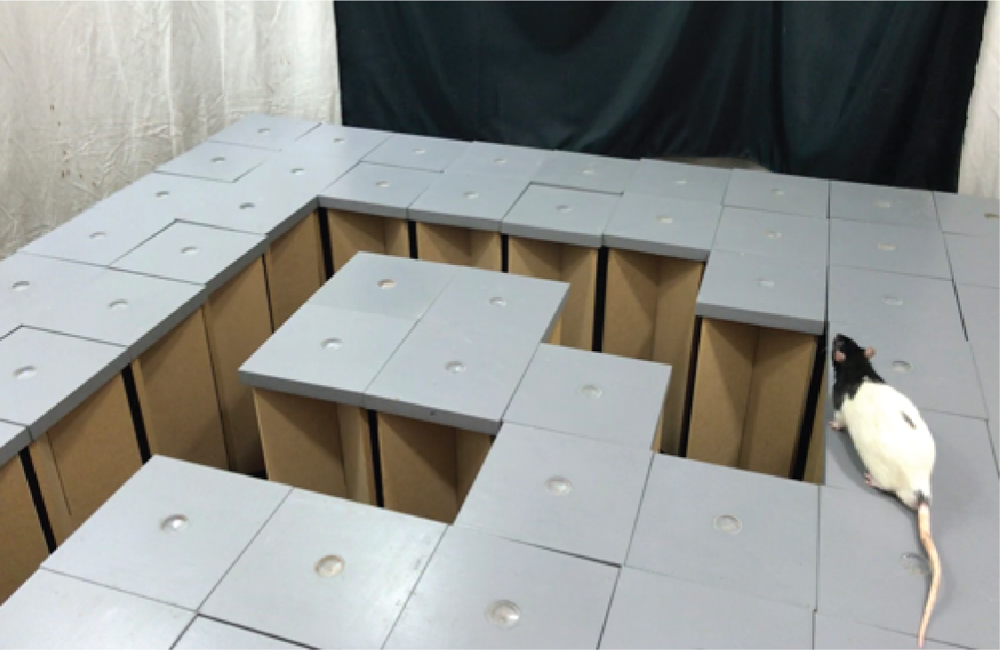
The maze environment used for the rat experiment. The environment consists of 100 removable mazes modules with a black curtain over one of the surrounding edges to provide a single extra-maze cue. Reward can be dispensed at the goal module by filling the well with chocolate milk via polymeric tubing beneath the maze.

**Figure S2:**
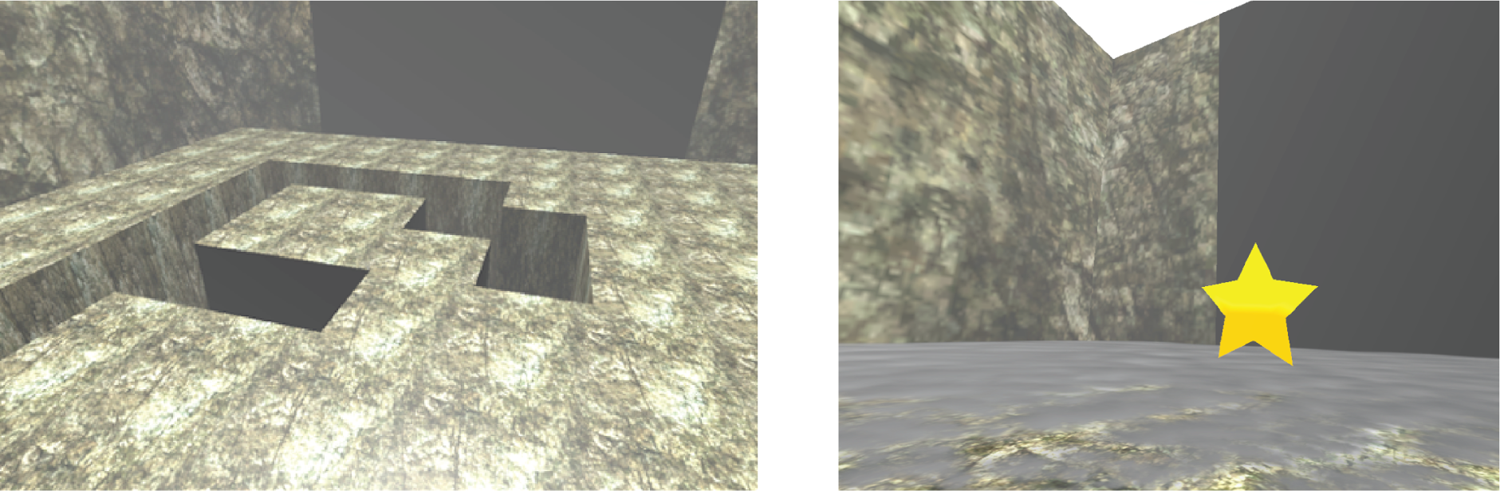
The virtual environment used for the human experiment. The environment had the same proportions as the rat environment and consisted of 100 removable mazes modules with a black curtain over one of the surrounding edges to provide a single extra-maze cue. A seamless texture was applied to the maze modules and walls and a fog lined the floor of the maze (see right image) to ensure humans had to rely on spatial memory to understand the maze structure. Reward was indicated by a gold star that would appear at the goal module when the participant successfully navigated to it.

**Figure S3:**
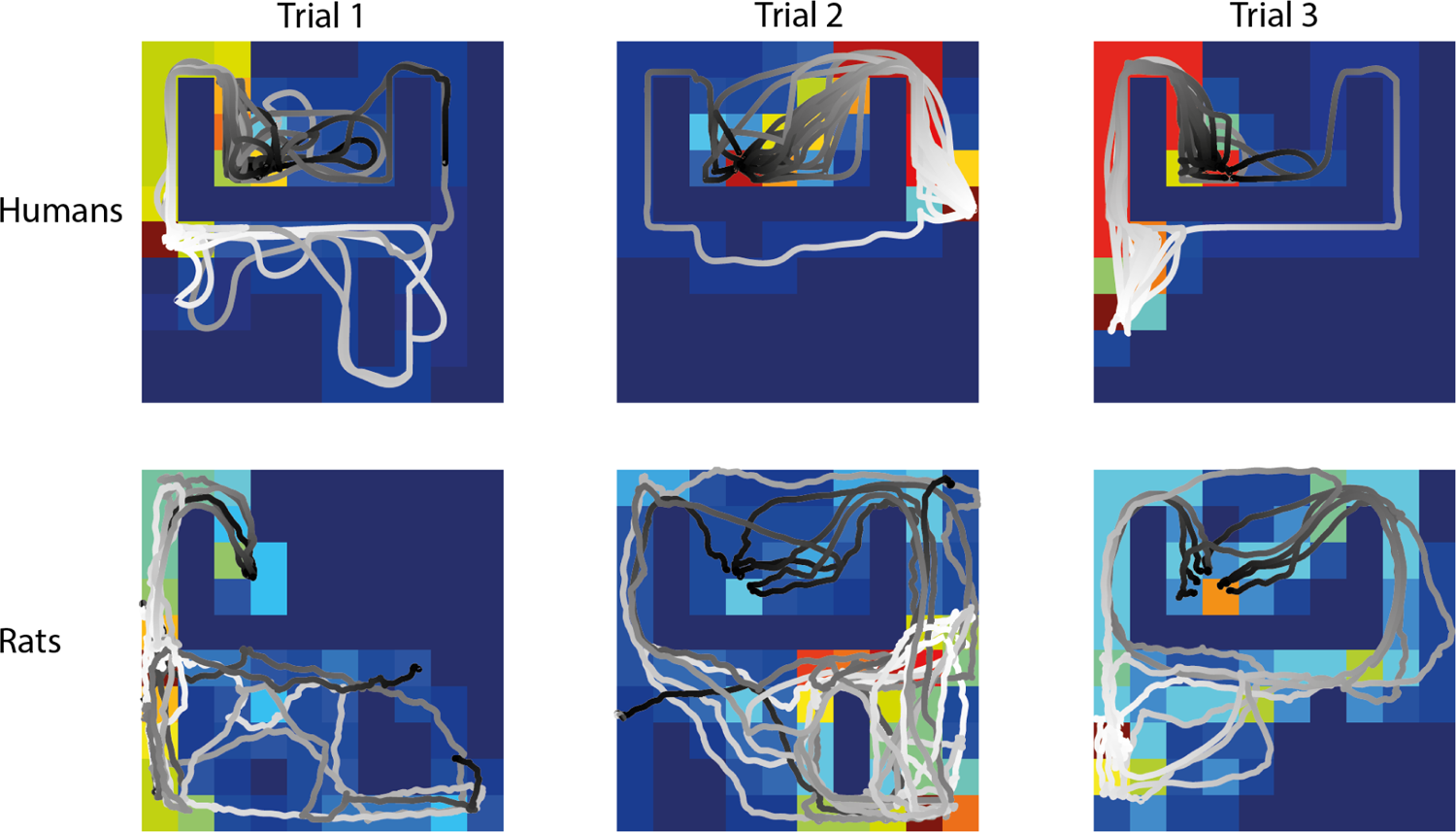
The route choices of humans and rats were often suboptimal at the start of a new maze configuration. Examples of the human (top) and rat (bottom) trajectories overlaying occupancy maps for the first 3 trials on maze configuration 20. The white-black colour gradient shows the beginning-end of each trajectory. Initially the paths taken by the humans and rats were often suboptimal (leftmost column) with performance generally improving rapidly within the first 3 trials of a new maze configuration. The goal location is 4 squares right and down from the top-left corner.

**Figure S4:**
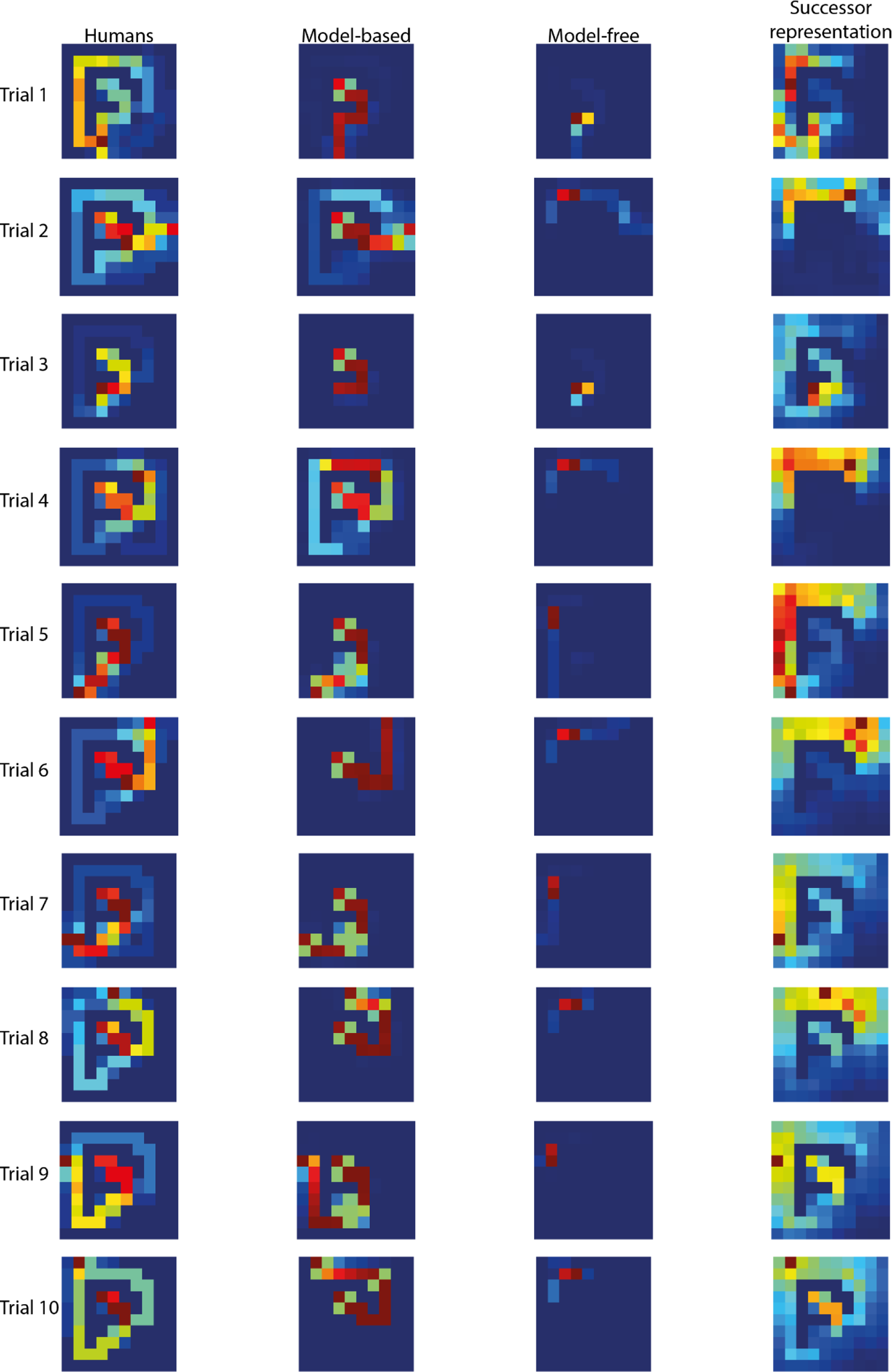
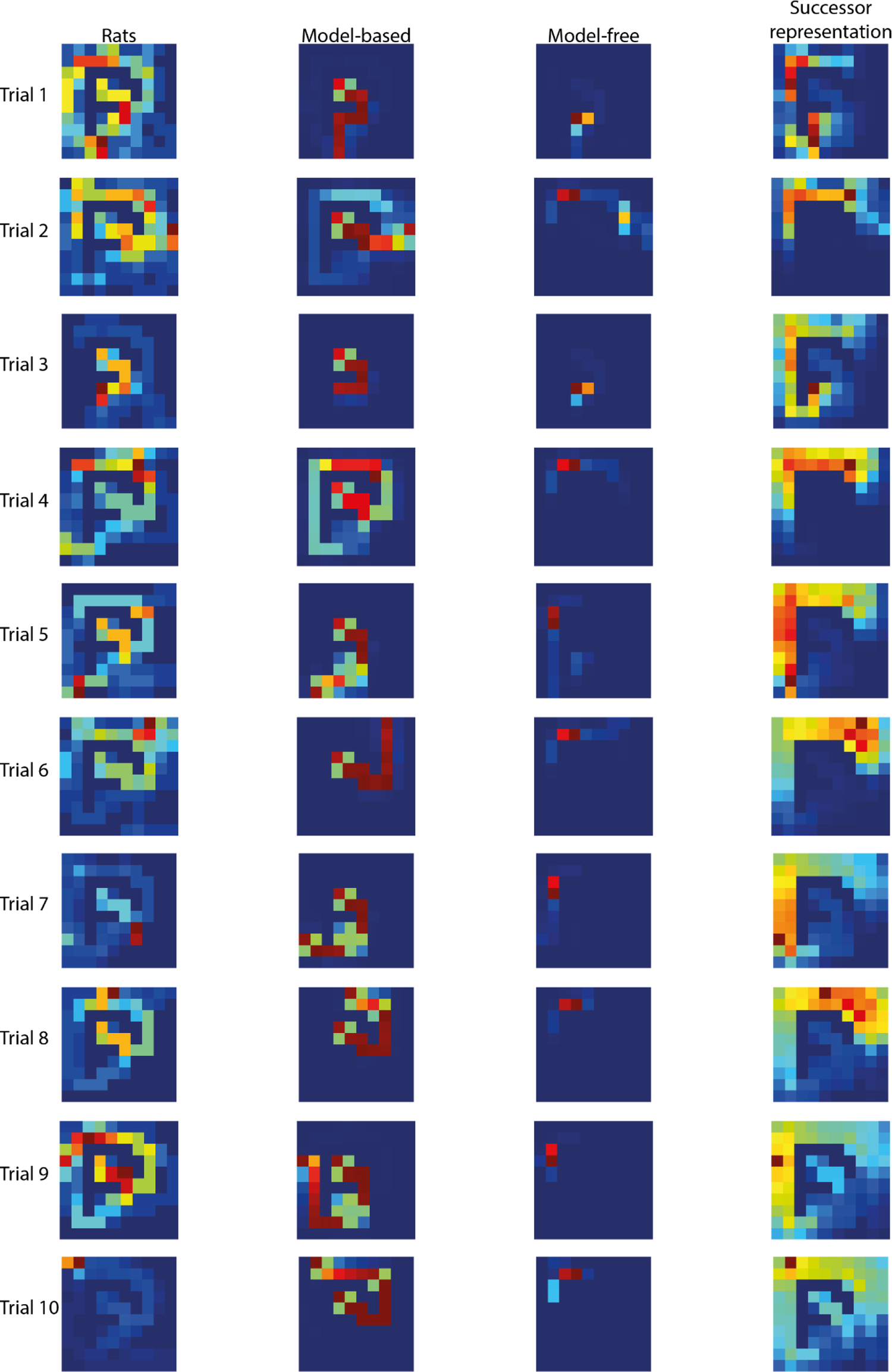
Human and agent occupancy maps for maze configuration 21. The occupancy maps of the humans (leftmost column) and agents for each of the 10 trials (rows) on maze configuration 21. The Model-Based agent (second column) quickly learns an accurate model of the environment and uses it to choose the shortest route to the goal with respect to that model (goal location is 4 squares right and down from the top-left corner). Conversely, the model-free agent (third column) is unable to update its value representation fast enough to successfully adapt to the new maze configuration, and particularly struggles on later trials where the starting position requires longer and more tortuous routes. The successor representation agent (rightmost column) sits on the spectrum between model-based and model-free methods, initially struggling to find an efficient route to the goal but providing a good match to the human behaviour on later trials.

**Figure S5:**
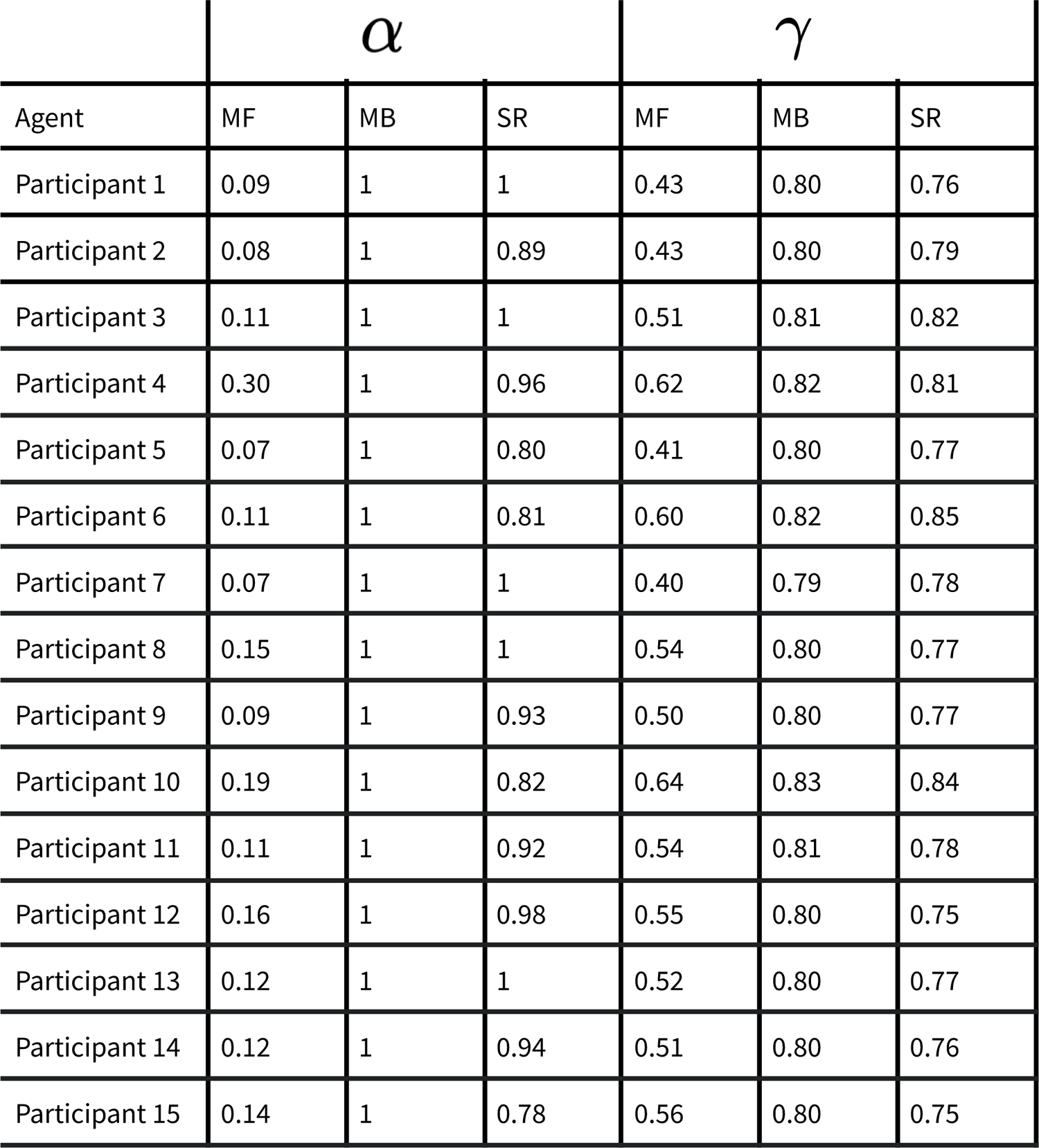
Rat and agent occupancy maps for maze configuration 21. The occupancy maps of the rats (leftmost column) and agents for each of the 10 trials (rows) on maze configuration 21. The Model-Based agent (second column) quickly learns an accurate model of the environment and uses it to choose the shortest route to the goal with respect to that model (goal location is 4 squares right and down from the top-left corner). Conversely, the model-free agent (third column) is unable to update its value representation fast enough to successfully adapt to the new maze configuration, and particularly struggles on later trials where the starting position requires longer and more tortuous routes. The successor representation agent (rightmost column) sits on the spectrum between model-based and model-free methods, initially struggling to find an efficient route to the goal but providing a good match to the rat behaviour on later trials.

---

**Table S1:**
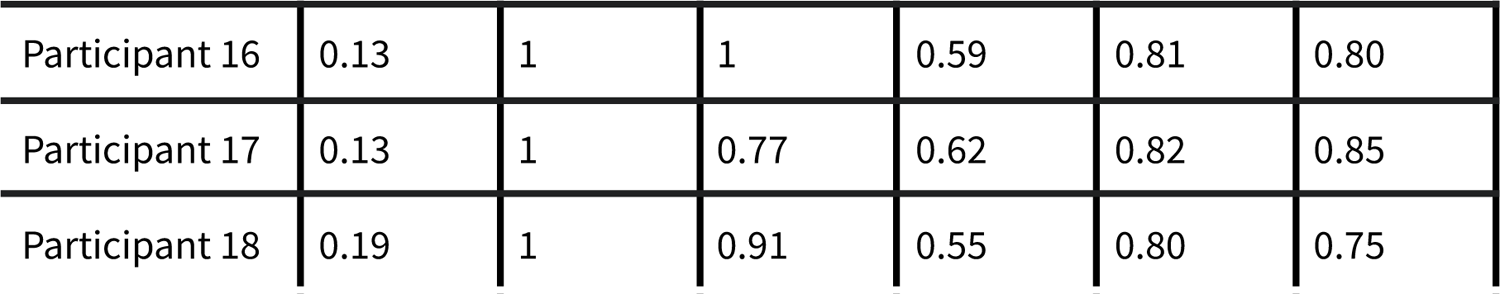
Human behaviour maximum likelihood parameters. The learning rates α and discount factors γ for the model-free (MF), model-based (MB) and successor representation (SR) agents, calculated for each individual.

**Table S2:** Rat behaviour maximum likelihood parameters. The learning rates α and discount factors γ for the model-free (MF), model-based (MB) and successor representation (SR) agents, calculated for each individual.

| | $\alpha$ | | | $\gamma$ | | |
| --- | --- | --- | --- | --- | --- | --- |
| Agent | MF | MB | SR | MF | MB | SR |
| Rat 1 | 0.07 | 1 | 0.76 | 0.43 | 0.79 | 0.80 |
| Rat 2 | 0.11 | 1 | 0.77 | 0.48 | 0.80 | 0.83 |
| Rat 3 | 0.12 | 1 | 0.87 | 0.45 | 0.79 | 0.71 |
| Rat 4 | 0.65 | 1 | 0.87 | 0.01 | 0.78 | 0.76 |
| Rat 5 | 0.42 | 1 | 0.77 | 0.01 | 0.78 | 0.77 |
| Rat 6 | 0.09 | 1 | 0.90 | 0.34 | 0.79 | 0.79 |
| Rat 7 | 0.12 | 1 | 0.70 | 0.16 | 0.80 | 0.82 |
| Rat 8 | 0.08 | 1 | 0.91 | 0.28 | 0.78 | 0.81 |
| Rat 9 | 0.07 | 1 | 0.87 | 0.42 | 0.80 | 0.81 |

**<u>Video S1</u>: Example rat trajectories**. An example of 10 trials (configuration 11) of Rat 2 from the rat version of the task. Video is sped up x2 after the first trial.

**<u>Video S2</u>: Example human trajectories**. An example of 10 trials (configuration 11) of participant 15 from the human version of the task. Video is sped up x2 after the first trial.

**<u>Video S3</u>: Example model-free agent trajectories.** Heatmap trajectories for 10 trials (configuration 11) of 100 model-free agents from the reinforcement learning simulation of the task, trained on participant 15. Agent density is represented by the shade of grey/black, with the green square indicating the goal location and the red squares indicating barriers in the environment.

**<u>Video S4</u>: Example model-based agent trajectories.** Heatmap trajectories for 10 trials (configuration 11) of 100 model-based agents from the reinforcement learning simulation of the task, trained on participant 15. Agent density is represented by the shade of grey/black, with the green square indicating the goal location and the red squares indicating barriers in the environment.

**<u>Video S5</u>: Example successor representation agent trajectories.** Heatmap trajectories for 10 trials (configuration 11) of 100 successor representation agents from the reinforcement learning simulation of the task, trained on participant 15. Agent density is represented by the shade of grey/black, with the green square indicating the goal location and the red squares indicating barriers in the environment.

